# Combined lactate and hypothermia therapy enhances neuroprotection following neonatal hypoxic-ischemic brain injury

**DOI:** 10.64898/2026.09.24.754268

**Authors:** Ifrah Omar Ibrahim, Sébastien Giraud, Estelle Lemarié, Marie-Christine Beauvieux, Stéphane Sanchez, Marc Biran, Aurélien Trottier, Karine Viaud, Jean-François Chateil, Jean Delmas, Luc Pellerin, Hélène Roumes, Anne-Karine Bouzier-Sore

## Abstract

**Background and Purpose:** Neonatal hypoxia-ischemia (NHI) is a leading cause of mortality and long-term neurodevelopmental disability. Therapeutic hypothermia (TH) is currently the only established treatment for NHI; however, its efficacy remains limited, underscoring the need for adjunctive neuroprotective strategies. Lactate is now regarded as a major metabolic substrate of the brain and has previously been shown to exert neuroprotective effects following experimental NHI. We investigated whether lactate enhances hypothermia-induced neuroprotection and evaluated the optimal timing of administration and potential underlying mechanisms.

**Methods:** Postnatal day 7 Wistar rats were subjected to NHI using the Rice-Vannucci model and treated with TH alone (2h at 32°C), sodium L-lactate alone, or combined L-lactate–TH. Animals underwent longitudinal magnetic resonance imaging (MRI) and behavioral assessments, complemented by histological, immunohistochemical, and Western blot analyses, including assessment of protein lactylation. The effect of sodium L-lactate administration timing relative to TH was also evaluated.

**Results:** Lactate significantly enhanced the neuroprotective effects of TH, particularly when administered before hypothermia, resulting in greater lesion recovery, reduced neuroinflammation and apoptotic cell death, preserved neuronal integrity, and improved long-term motor and cognitive outcomes. Combined treatment preserved neuronal lactate transporter, prevented NHI-induced astrocytic glutamate transporter upregulation, modulated lactylation markers, supporting coordinated metabolic and epigenetic adaptations following NHI.

**Conclusions:** Early lactate administration enhances the neuroprotective efficacy of TH following NHI. These findings identify lactate as a promising adjunct to TH and support its translational potential as a metabolic strategy to improve outcomes following NHI brain injury.

## Introduction

Neonatal hypoxia-ischemia (NHI) remains a leading cause of neonatal mortality and long-term neurodevelopmental impairment worldwide, affecting approximately 0.7 to 1.2 million newborns annually and accounting for nearly 23% of neonatal deaths.^1^ Its incidence ranges from 1 to 6 per 1,000 live births in high-income countries to 26.5 per 1,000 in low- and middle-income regions. Despite advances in neonatal care, 15–20% of affected infants die during the neonatal period, and up to 25% of survivors develop severe neurological sequelae, including cerebral palsy, epilepsy, and cognitive impairment.^2^

Therapeutic hypothermia (TH) is currently the only approved treatment for NHI. Initiated within 6h of injury and maintained for 72h, TH provides neuroprotection and improves outcomes after NHI.^3,4^ Despite these benefits, 44–53% of treated infants either die or survive with moderate-to-severe disability,^5^ underscoring the urgent need for new therapy. Accordingly, numerous pharmacological and metabolic strategies have been evaluated as adjuncts to TH in preclinical models, including rodent and piglet models of NHI.^6–11^ Although several strategies have demonstrated promising neuroprotective effects and some have progressed to clinical evaluation, no adjunct therapy has yet been approved for clinical use in combination with TH.

Once considered merely a metabolic waste product, lactate is now recognized as a metabolic substrate of the brain, and particularly of neurons.^12,13^ According to the astrocyte-neuron lactate shuttle (ANLS) concept, astrocytes produce lactate via aerobic glycolysis and transfer it to neurons through monocarboxylate transporters (MCTs), where it is oxidized to support neuronal energy demands.^14–16^ The functional relevance of the ANLS has been supported by data obtained *in vivo*, pointing out the role of lactate as an important metabolic substrate for neuronal function and memory.^17–19^ Lactate availability may therefore sustain neuronal energy metabolism when glucose availability or utilization is limited.

Consistent with this concept, lactate administration has emerged as a promising neuroprotective strategy in conditions associated with cerebral energy failure, including experimental ischemic stroke and traumatic brain injury.^20–23^ This rationale is particularly relevant to the neonatal brain, which is highly adapted to the utilization of alternative monocarboxylate fuels. During the suckling period, ketone bodies and lactate readily cross the blood-brain barrier via highly expressed MCTs, while glucose transporter expression is comparatively lower.^24,25^ This metabolic adaptation favors the use of lactate as an alternative substrate for neurons in neonates.

In the context of NHI, several studies have demonstrated that lactate administration confers neuroprotection.^26,27^ Beyond its metabolic role, lactate has also been shown to act as a signaling molecule in experimental hypoxia-ischemia models, promoting angiogenesis and contributing to microvascular repair.^28,29^ However, whether the neuroprotective effects of lactate previously observed in NHI are preserved when lactate is administered in combination with TH, the current standard of care for moderate-to-severe NHI, remains unknown. Here, we evaluated the neuroprotective potential of sodium L-lactate administered within the therapeutic window in combination with TH in a rat model of NHI. Using a longitudinal design combining MRI and behavioral assessments, we assessed structural and functional outcomes, and also explored the molecular mechanisms underlying the neuroprotection with additional histological and protein analyses.

## Materials and Methods

### Animals

All animal procedures were conducted in accordance with the European Communities Council Directive (86/609/EEC, 24 November 1986), the ethical guidelines of the French Ministry of Agriculture and Forestry, and the ARRIVE guidelines. Experimental protocols were approved by the Institutional Animal Care and Use Committee of the University of Bordeaux (C2EA-50; authorization no. 37955).

Timed-pregnant female Wistar RJHan rats (Janvier, France) were received on gestational day 16 and housed under standard laboratory conditions (12h light/12h dark cycle, ambient temperature 21– 22°C, relative humidity 55–60%) with *ad libitum* access to food and water. Animals were housed in ventilated polycarbonate cages with wood-chip bedding. Litters were standardized to a maximum of eight pups of either sex. Every effort was made to minimize animal suffering and to use the minimum number of animals.

### Model of Neonatal Hypoxia-Ischemia

NHI was induced at postnatal day 7 (P7) using a modified Rice-Vannucci model,^30^ as previously described in our laboratory.^26,31,32^ Briefly, pups were anesthetized with isoflurane, and the left common carotid artery was permanently ligated. Following a 30-min recovery period, animals were exposed to a humidified gas mixture containing 8% O₂ and 92% N₂ for 2h in a temperature-controlled chamber (35 ± 1°C). Sham-operated animals underwent identical surgical procedures without carotid artery ligation and were maintained under identical normothermic conditions.

### Experimental groups

The experimental design is summarized in **Fig.1**. Animals were assigned to the following six experimental groups: **Sham**: Surgery without occlusion; saline, normothermia (2h); **HIC**: HI+saline; normothermia (2h); **HIL**: HI+lactate; normothermia (2h); lactate at 24 and 48h; **HIH**: HI+hypothermia (2h); **HIL-H**: HI+lactate immediately before hypothermia (2h); lactate at 24 and 48h and **HIH-L**: HI+hypothermia (2h), followed by lactate; lactate at 24 and 48h; MRI/behavior only.

**Figure. 1.**
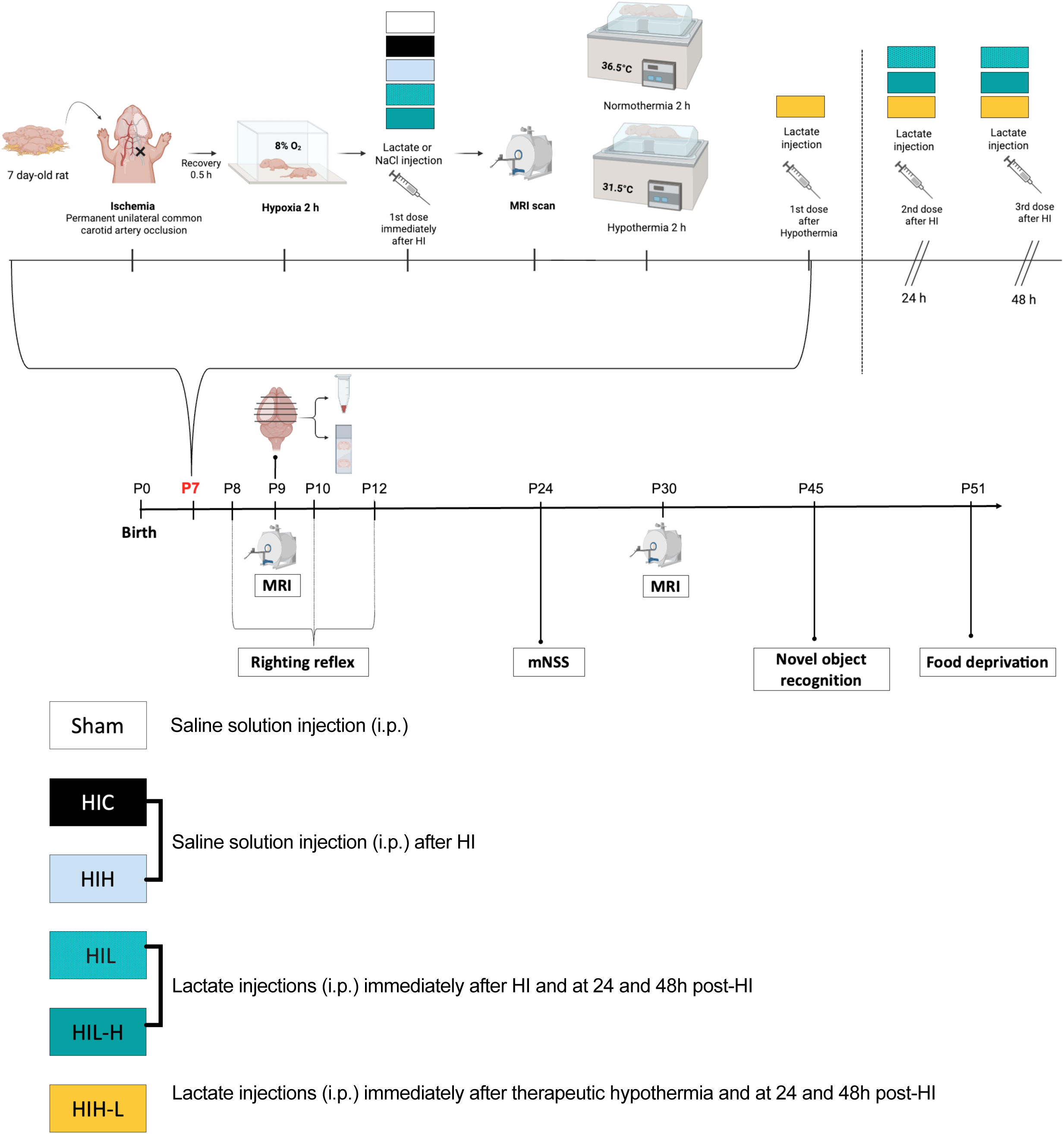
Experimental design and study groups. A total of 135 postnatal day 7 (P7) Wistar rat pups were initially included in the study. Seven animals died during or immediately after the hypoxic insult, resulting in 128 animals available for analysis. Of these, 49 animals were euthanized at P9 for histological, immunohistochemical, and Western blot analyses, whereas the remaining 55 animals underwent longitudinal MRI and behavioral assessments. Animals were allocated to the following experimental groups: Sham (n = 13; 8 males, 5 females), HIC (n = 18; 8 males, 10 females), HI+lactate (HIL) (n = 12; 7 males, 5 females), HI+hypothermia (HIH) (n = 11; 5 males, 6 females), HI+lactate+hypothermia (HIL-H) (n = 13; 5 males, 8 females), and HI+hypothermia+lactate (HIH-L) (n = 12; 6 males, 6 females). Except for the Sham group, all animals underwent neonatal hypoxia-ischemia (NHI). Animals in the HIL and HIL-H groups received sodium L-lactate immediately after hypoxia and again at 24 and 48h post-HI. Animals in the HIH-L group received the first lactate injection after TH, followed by additional injections at 24 and 48h post-NHI. The longitudinal evolution of brain lesion volume was assessed MRI at P7, P9, and P30. Behavioral assessments included the righting reflex (P8, P10, and P12), the mNSS (P24), the NOR (P45), and the food restriction behavioral tests (P51).

As 2h was previously identified as the optimal duration of TH in this NHI rat model,^326,337^ here we evaluated whether lactate provides additional neuroprotection when administered in combination with this optimized hypothermia protocol. Forty-nine animals were euthanized at P9 for *ex vivo* analyses, whereas the remaining animals underwent longitudinal MRI and behavioral assessments. Pups from each litter were randomly allocated among the experimental groups.

### Hypothermia and normothermia treatment

TH was performed as previously established in our laboratory.^32^ Briefly, animals assigned to HIH, HIL-H and HIH-L underwent 2h of hypothermia initiated 3.5h after carotid ligation. Core temperature was maintained at 32.0 ± 0.5°C and pups were progressively rewarmed before being returned to their dams. Sham, HIC, and HIL animals were maintained under normothermic conditions for the same duration using the previously established protocol.^36^

### Lactate administration

Animals received three intraperitoneal injections of sodium L-lactate (AP-HP 11.2% w/v; 999 mmol/L sodium; 999 mmol/L lactate; 4 µL/g body weight). The administered dose was (4.2 mmol/kg/day for 3 consecutive days). In the HIL and HIL-H groups, sodium L-lactate was administered immediately after hypoxia (0h) and at 24 and 48h post-HI. Sham and HIC animals received equivalent volumes of 0.9% NaCl at the same time points. In the HIH-L group, the same dosing schedule was followed, except that the first lactate injection was administered after the 2h hypothermia period (3h post-HI). For simplicity, sodium L-lactate is referred to as “lactate” throughout the manuscript

### In vivo MRI

MRI acquisition and lesion quantification were performed longitudinally in the same animals as previously described.^32^ Briefly, MRI was performed at P7, P9, and P30 using a 4.7T Bruker Biospec system. Diffusion-weighted imaging (DWI) was acquired at P7 and P9 to assess acute brain injury, whereas T2-weighted anatomical imaging was performed at P30 to evaluate long-term lesion outcome. Lesion volumes were quantified from apparent diffusion coefficient (ADC) maps at P7 and P9 and from T2-weighted images at P30. Regions of interest corresponding to injured tissue were manually delineated and expressed as a percentage of total brain volume. Detailed MRI acquisition parameters and image processing procedures are provided in the Supplemental Methods.

### Behavior

Cognitive, sensory, and motor functions were assessed between P8 and P51 (spanning a developmental period roughly corresponding to early infancy through adolescence in humans), as previously described.^32^

#### Righting reflex

Motor coordination was assessed at P8, P10 and P12 by recording the latency to regain the prone position from supine, averaged over three trials.

#### Modified neurological severity score (mNSS)

Neurological function was assessed at P24 using the mNSS (0–18), which evaluates through different tasks motor function, sensory responses, reflexes, and balance. Each failed task scores points, with higher scores indicating greater impairment.

#### Novel object recognition

Recognition memory was assessed from P42 to P45. Following habituation (1 day) and familiarization with two identical objects (2 days), one was replaced by a novel object (trial day). Exploration was recorded for 5min, and memory performance was quantified using the discrimination index [(novel object exploration time – familiar object exploration time)/ (novel object exploration time + familiar object exploration time)], recognition index [novel object exploration time/ (novel object exploration time + familiar object exploration time)], and total exploration time.

#### Food deprivation test

Stress-related and feeding behavior was assessed at P48–P51 following progressive food restriction over 3 days.^36^ Exploratory behavior, food investigation, latency to pellet consumption, and bruxism were recorded.

### Immunohistochemistry

Three to five animals per group were selected at P9 based on their P7 MRI lesion volumes being within the average range of their respective groups, ensuring representative sampling while maximizing the number of animals available for longitudinal imaging and behavioral assessments. Anesthetized animals were then perfused with PBS followed by 4% paraformaldehyde, and their brains were processed for cryostat sectioning as previously described.^32^ Immunohistochemical analyses were performed on 16-µm cryostat sections from the ipsilateral cortex and hippocampus. Sections were stained with antibodies against cleaved caspase-3 and Iba1 to assess apoptotic cell death and microglial activation, respectively. Images were acquired using a Nikon Eclipse 90i microscope, and quantitative analyses were performed by an investigator blinded to experimental groups. Detailed immunostaining procedures and additional microglial polarization analyses are provided in the Supplemental Methods.

### Western blots

Ipsilateral and contralateral cortical tissues were collected (5 to 7 P9 animals/group, selected as described above), snap-frozen, and processed for protein extraction. Protein expression was assessed by SDS-PAGE and immunoblotting using antibodies listed in **Table S2**. Signals were quantified and normalized to total protein staining or β-actin or GAPDH, depending on the targeted protein, and expressed relative to the Sham group, set at 100%.

### Statistical analysis

Except for bruxism and the modified neurological severity score (mNSS), data are presented as mean ± SEM. Bruxism data are reported as the mean with range, whereas mNSS data are presented as median (interquartile range [IQR]). Statistical analyses were performed using GraphPad Prism version 10. Data normality was assessed using the Shapiro-Wilk test. For normally distributed datasets, one-way analysis of variance (ANOVA) was used for multiple-group comparisons, followed by Tukey’s or Fisher’s post hoc multiple-comparisons test, as specified in the corresponding figure legends. For datasets that did not meet normality assumptions, the Kruskal-Wallis’ test was performed, followed by Dunn’s multiple-comparisons test. Pairwise comparisons between the HIL-H and HIH-L groups, as well as sex-based comparisons within each experimental group, were performed using an unpaired two-tailed Student’s t test or, when normality assumptions were not met, the Mann-Whitney U test. One outlier was identified using the ROUT method (Q = 1%) only for ipsilateral H3K9la analyse. A two-sided P < 0.05 was considered statistically significant.

## Results

### Lactate administration reduces early brain injury after NHI

To confirm that lactate provides early neuroprotection after NHI, DWI was performed 3h after carotid artery ligation and 30min after intraperitoneal injection of lactate (or saline). Representative images and lesion quantification are shown in **Fig.2A–B**. HI induced extensive diffusion abnormalities consistent with cytotoxic edema, whereas lactate-treated pups exhibited a marked reduction in early lesion burden. Lesion volume was significantly decreased in the HIL group compared with untreated HIC animals (34 ± 1% *vs.* 46 ± 1% of total brain volume; respectively; **Fig.2B**). Consistent with reduced tissue injury, lactate administration attenuated HI-induced decreases in ADC values across vulnerable brain regions. Compared with sham animals, HIC pups displayed pronounced ADC reductions in the cortex (-44%), hippocampus (-42%), and striatum (-37%), reflecting restricted water diffusion associated with cytotoxic edema. Lactate treatment attenuated the decrease in ADC values in the cortex (-34%) and hippocampus (-25%) and restored ADC values in the striatum. Across all analyzed regions, ADC values remained significantly higher in HIL pups than in HIC animals (**Fig.2C**). Together, these findings demonstrate that the initial lactate administration, given immediately after NHI as part of the three-dose treatment regimen, reduces lesion volume and attenuates cytotoxic edema at this early time point.

**Figure. 2.**
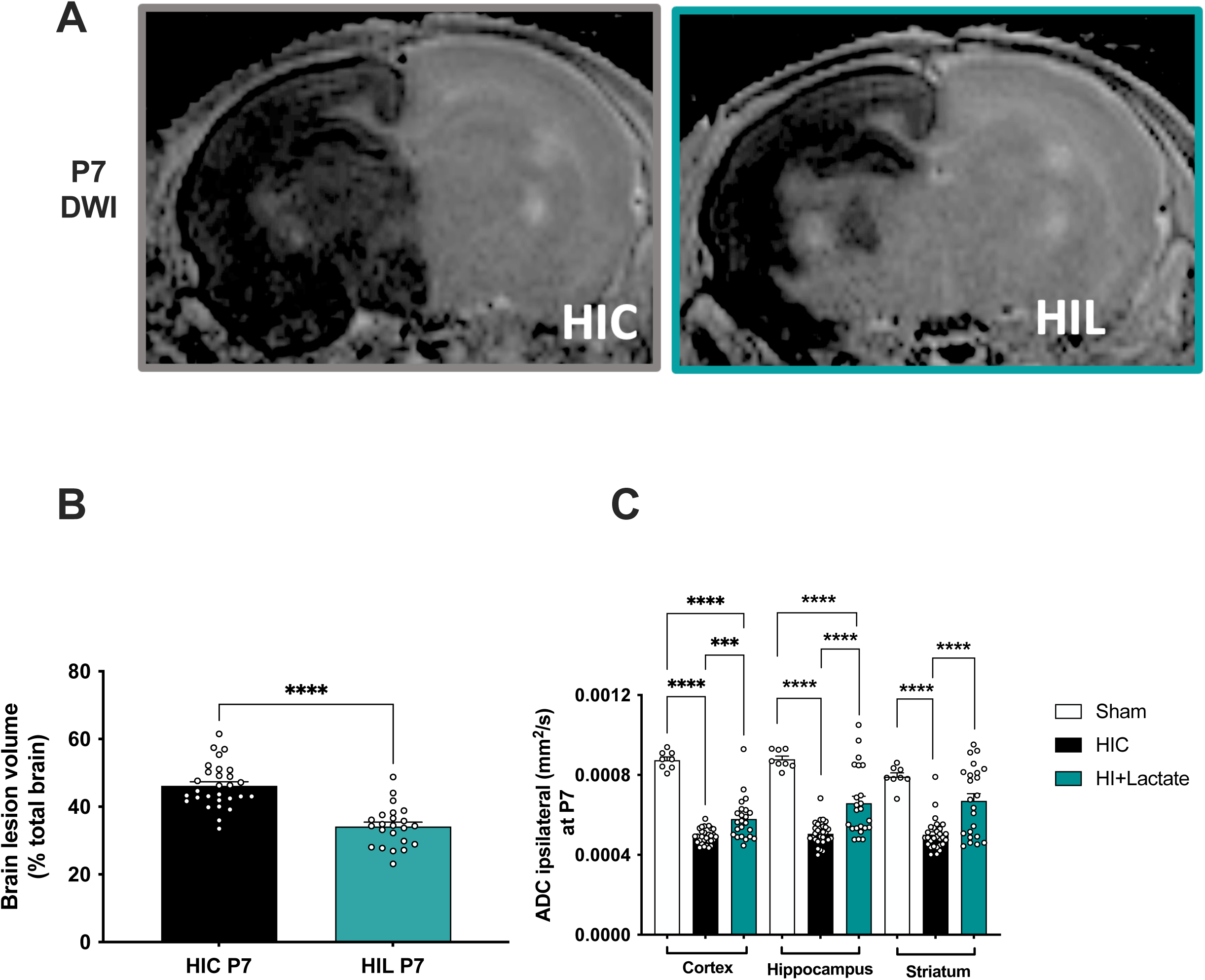
Early MRI assessment of brain injury following NHI and lactate administration. (A) Representative apparent diffusion coefficient (ADC) maps of P7-rat brains from HIC and HI+Lactate groups acquired 3h after carotid artery ligation using a diffusion-weighted imaging (DWI) sequence. Brain injury is visualized as a hypointense signal on ADC maps, reflecting reduced water diffusivity associated with cytotoxic edema. (B) Quantification of brain lesion volume identified from ADC maps and expressed as a percentage of total brain volume. (C) Quantification of ipsilateral ADC values in the cortex, hippocampus, and striatum from Sham (n = 8), HIC (n = 29), and HI+Lactate (n = 23) animals. Data are presented as mean ± SEM. Statistical analyses were performed using an unpaired Student’s t-test or one-way ANOVA followed by Tukey’s multiple-comparisons test, as appropriate. Statistical significance is indicated as ***p < 0.001 and ****p < 0.0001.

### Combined lactate and hypothermia treatment enhances early lesion recovery following NHI

We next investigated whether lactate retains its neuroprotective effects when combined with TH, by comparing the combined treatment with lactate or TH alone. Brain lesion volumes were longitudinally assessed by MRI (**Fig.3A**). At P9, both lactate treatment alone and combined lactate-TH treatment markedly reduced lesion volumes compared with HIC animals. Lesion volumes were 18 ± 2% in the HIL-H group, 21 ± 2% in the HIL group, 29 ± 2% in the HIH group, and 38 ± 1% in the HIC group (**Fig.3B**). Lesion volumes did not differ significantly between HIL and HIL-H groups, although combined treatment resulted in significantly smaller lesions than TH alone, indicating enhanced early tissue protection.

**Figure. 3.**
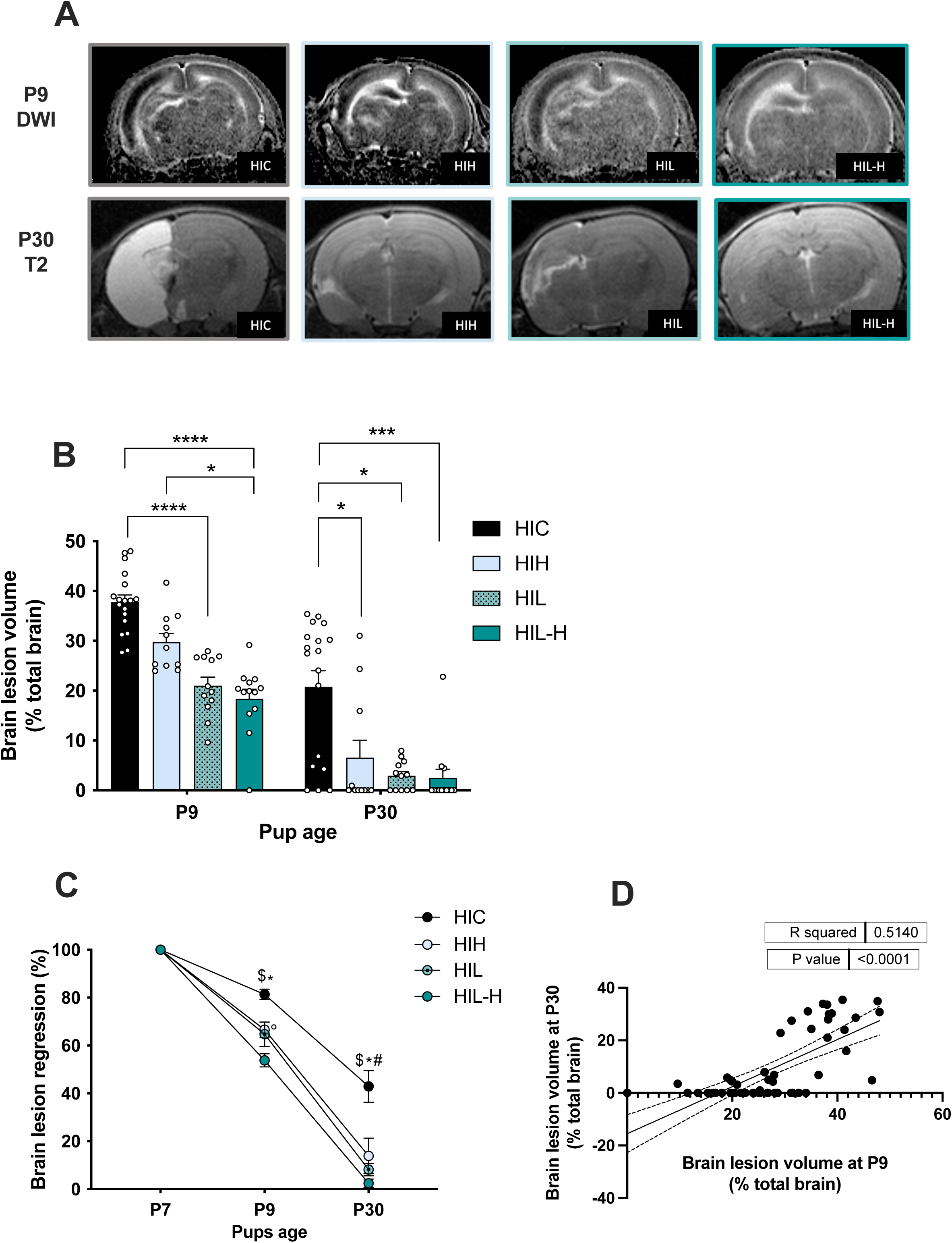
Longitudinal MRI assessment of brain injury following NHI. (A) Representative apparent diffusion coefficient (ADC) maps derived from diffusion-weighted imaging (DWI) at P7 (initial injury), P9, and T2-weighted images acquired at P30 from animals in the HIC (n = 18), HIH (n = 11), HIL (n = 12), and HIL-H (n = 13) groups using a 4.7 T MRI system. Lesions appear as hypointense signal on ADC maps at P7 and P9 and as hyperintense signal on T2-weighted images at P30. (B) Quantification of lesion volumes expressed as a percentage of total brain volume across experimental groups at P7, P9, and P30. (C) Quantification of brain lesion regression over time, expressed as the percentage of lesion volume reduction relative to the initial lesion volume measured at P7. (D) Correlation between brain lesion volumes measured by MRI at P9, and P30. Data are presented as mean ± SEM. Statistical analyses were performed using Kruskal-Wallis tests followed by Dunn’s multiple-comparisons test. Statistical significance is indicated as *p < 0.05, ***p < 0.001, and ****p < 0.0001. In panel (C), * indicates comparisons between HIC and HIL, $ between HIC and HIL-H, ° between HIH and HIL-H, and # between HIC and HIH.

At P30, brain lesion volumes remained significantly lower in all treated groups than in HIC animals (HIL-H: 2 ± 2%; HIL: 3 ± 1%; HIH: 7 ± 3%; HIC: 21 ± 3%; **Fig.3B**). However, no significant differences were observed among the HIL-H, HIL, and HIH groups, suggesting that all three treatments provided comparable long-term structural recovery.

Analysis of brain lesion evolution from P7 to P30 revealed accelerated early recovery in lactate-treated animals. Within the first 48h after NHI, lesion recovery reached 35 ± 5% in the HIL group and 51 ± 5% in the HIL-H group, with the combined treatment showing the highest early recovery rate (**Fig.3C**). By P30, all treated groups showed substantial lesion resolution compared with HIC animals.

Finally, correlation analyses demonstrated a significant association between brain lesion volumes measured at P9 and P30, indicating that early lesion burden was predictive of subsequent lesion outcome at the individual-animal level (**Fig.3D**).

### Combined lactate and hypothermia treatment improves long-term motor and cognitive recovery following NHI

To determine whether the structural improvements observed by MRI were associated with functional recovery, we next assessed motor and cognitive outcomes using longitudinal behavioral testing.

Motor coordination was evaluated from P8 to P12 using the righting reflex test. HIC animals displayed persistent motor impairment throughout the assessment period, with no improvement over time compared with sham animals (**Fig.4A**). In contrast, lactate-treated animals exhibited rapid functional recovered, with HIL pups performing similarly to sham animals at all time points. HIH and HIL-H animals showed intermediate recovery with improved performance compared with HIC animals but residual impairment relative to Sham and HIL groups during the early assessment period. By P12, HIL-H animals achieved complete recovery of the righting reflex, whereas HIH animals showed only partial recovery (**Fig.4A**).

**Figure. 4.**
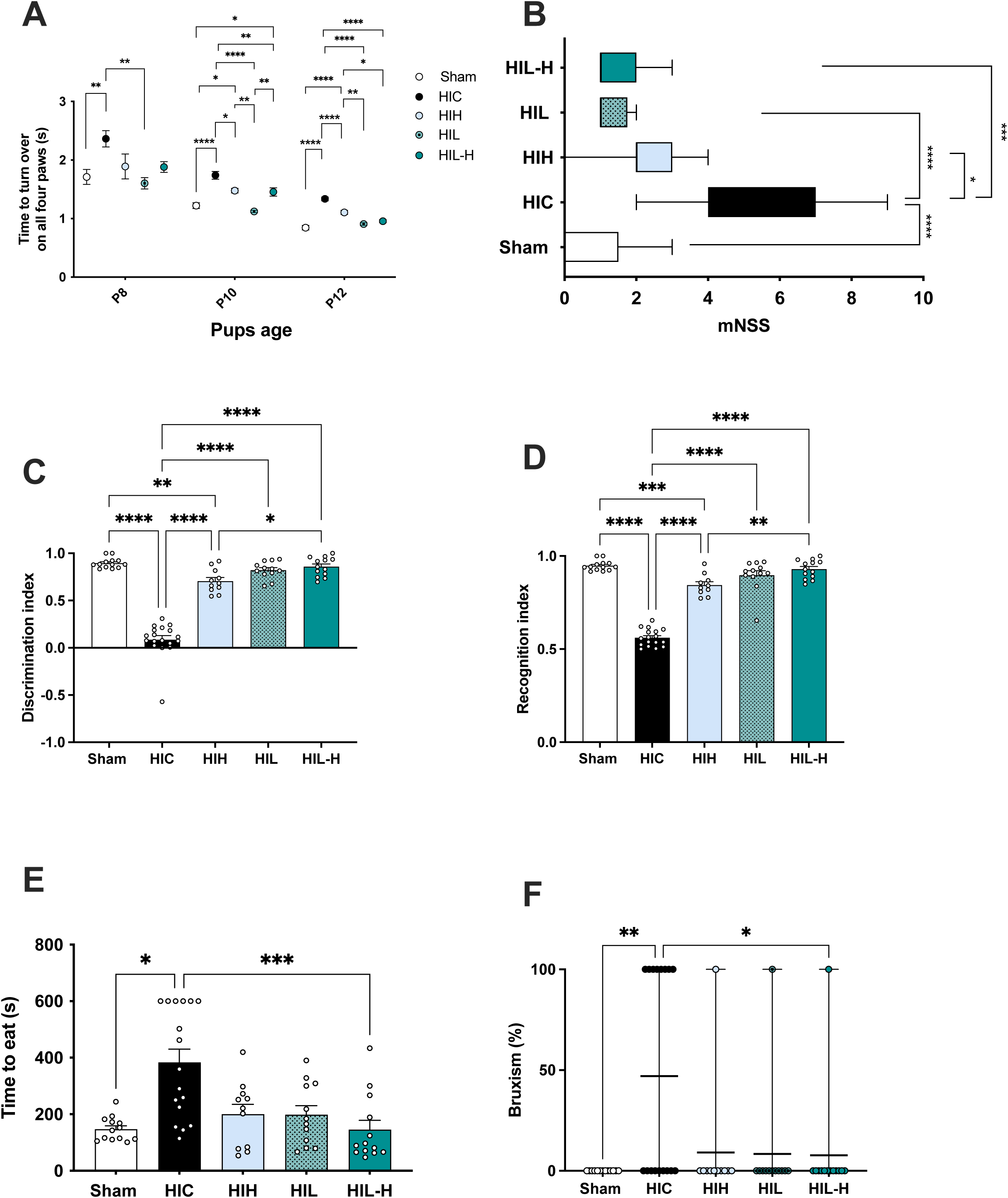
Longitudinal behavioral assessment. (A) Righting reflex assay assessed at P8, P10, and P12 in Sham (n = 13), HIC (n = 18), HIH (n = 11), HIL (n = 12), and HIL-H (n = 13) groups. (B) Modified Neurological Severity Score (mNSS) evaluated at P24 across experimental groups. (C, D) Novel object recognition test performed at P45 showing the discrimination index (C) and recognition index (D). (E, F) Food restriction test performed at P51 measuring the latency to consume the food pellet (E) and the percentage of animals displaying bruxism (F) in Sham (n = 13), HIC (n = 17), HIH (n = 11), HIL (n = 12), and HIL-H (n = 13) groups. Data are presented as mean ± SEM, except for bruxism data, which are presented as mean with range, and mNSS data, which are presented as median with interquartile range (IQR). Statistical analyses were performed using ordinary one-way ANOVA followed by Tukey’s multiple-comparisons test for normally distributed data (A, C, D) or Kruskal-Wallis tests followed by Dunn’s multiple-comparisons test for non-normally distributed data (B, E, F). Statistical significance is indicated as *p < 0.05, **p < 0.01, ***p < 0.001, and ****p < 0.0001.

Sensorimotor deficits were further assessed at P24 using mNSS. HIC animals exhibited significantly greater neurological impairment compared with Sham and treated groups (mean scores: Sham, 0.8 ± 0.3; HIC, 5.5 ± 0.4; HIH, 2.2 ± 0.4; HIL, 1.3 ± 0.1; HIL-H, 1.5 ± 0.2; **Fig.4B**). No significant differences were observed among HIH, HIL, HIL-H, and sham animals, indicating substantial recovery following therapeutic intervention.

Because NHI is associated with persistent cognitive deficits, recognition memory was evaluated at P45 using the novel object recognition test. HIC animals displayed impaired memory performance, with significantly reduced discrimination and recognition indices compared with Sham (discrimination index: 0.09 ± 0.04 vs. 0.90 ± 0.01, respectively; **Fig.4C**). In contrast, TH alone, lactate alone, and combined lactate-TH treatment significantly improved discrimination performance (0.71 ± 0.04, 0.82 ± 0.03, and 0.86 ± 0.03, respectively). Notably, both lactate-treated groups, either alone or in combination with TH, achieved performance comparable to Sham, whereas the HIH group remained significantly different from Sham (**Fig.4C**). Similar findings were observed for the recognition index. HIC animals showed persistent impairment (0.56 ± 0.01), whereas HIL and HIL-H animals displayed recognition performance comparable to Sham (0.89 ± 0.02, 0.93 ± 0.01, and 0.95 ± 0.00, respectively; **Fig.4D**). HIH animals showed partial improvement (0.84 ± 0.02), while HIL-H animals exhibited significantly higher recognition performance than HIH animals (**Fig.4D**). No significant differences in the exploration index were observed among the Sham, HIH, HIL, and HIL-H groups compared with the HIC group (**Fig.S2A**).

Finally, emotional-like behaviors were assessed at P51 using the food restriction paradigm. HIC animals exhibited prolonged latency to consume food compared with sham animals, whereas HIH, HIL, and HIL-H groups showed latencies comparable to Sham (**Fig.4E**). In addition, bruxism was not observed in Sham (0%) but was present in 47% of HIC animals and was rarely detected in treated groups (9% in HIH, 8% in HIL, and 7% in HIL-H; **Fig.4F**), suggesting prevention of HI-induced stress-related behavioral alterations.

Correlation analyses revealed no significant associations between brain lesion volumes and behavioral outcomes, except in the HIC group, where P9 lesion volume correlated with the discrimination index (**Fig.S2B**).

### Sex-dependent analysis

No sex-related differences were detected in any experimental group across the previous assessed outcomes, including MRI measurements at P7, P9, and P30 and all behavioral tests performed throughout the study (**Fig.S3–S8**).

### Neuronal apoptosis following NHI

To determine whether the structural and functional improvements induced by lactate treatment were associated with reduced neuronal cell death, cleaved caspase-3 immunoreactivity was quantified in the ipsilateral cortex and hippocampal CA1 region 48h after NHI (**Fig.5A–C**). NHI markedly increased cleaved caspase-3 expression compared with sham animals in both the cortex (943 ± 40% *vs.* 100 ± 23%) and CA1 (496 ± 46% *vs.* 100 ± 12%; **Fig.5B–C**). Lactate treatment, either alone or combined with TH, significantly reduced cleaved caspase-3 immunoreactivity in both regions. TH alone also attenuated apoptotic cell death compared with HIC animals (**Fig.5B–C**). Consistent with these findings, Nissl staining confirmed improved neuronal preservation in treated animals, with the greatest preservation in the hippocampal CA1 region observed following combined lactate and TH treatment (**Fig.S9A–C**).

**Figure. 5.**
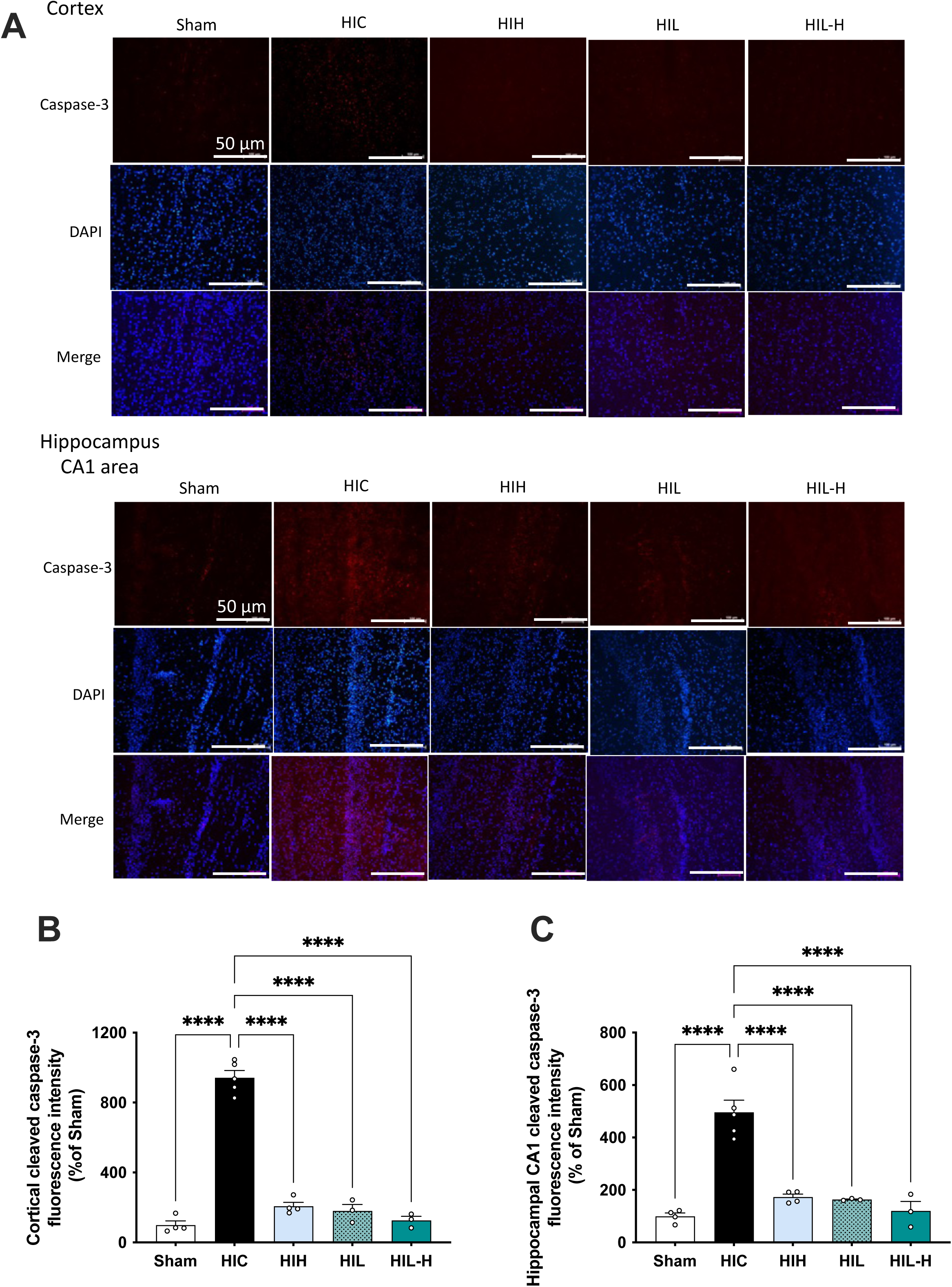
Early assessment of apoptotic cell death. (A) Representative immunofluorescence images showing cleaved caspase-3 immunoreactivity in the ipsilateral cortex and hippocampal CA1 region from Sham (n = 4), HIC (n = 5), HIH (n = 4), HIL (n = 3), and HIL-H (n = 3) groups at 48h after NHI (P9). Images were acquired at 20× magnification. Scale bar, 50 μm. (B, C) Quantification of cleaved caspase-3 immunoreactivity in the cortex (B) and hippocampal CA1 region (C). For each animal, fluorescence intensity was averaged from two independent fields per brain region. Data are presented as mean ± SEM. Statistical analyses were performed using ordinary one-way ANOVA followed by Tukey’s multiple-comparisons test. Statistical comparisons were performed relative to the Sham group. Significance is indicated as ****p < 0.0001.

### Microglial activation following NHI

Microglial activation was assessed by Iba1 immunostaining in the ipsilateral cortex and hippocampal CA1 region 48h after NHI (**Fig.6A–C**). NHI markedly increased Iba1 immunoreactivity compared with Sham in both regions (cortex: 722 ± 53% *vs.* 100 ± 8%; CA1: 875 ± 31% *vs.* 100 ± 22% for HIC and Sham groups, respectively; **Fig.6B–C**). Lactate treatment, alone or combined with TH, significantly reduced microglial activation in both regions. Although TH alone also decreased Iba1 immunoreactivity, combined treatment produced a greater reduction in the hippocampal CA1 region (148 ± 14% *vs.* 437 ± 76% HIL-H and HIH groups, respectively; **Fig.6C**). Consistent with these findings, all treated groups showed reduced CD86 expression and increased CD206 expression, supporting a shift toward a less pro-inflammatory microglial phenotype (**Fig.S10**).

**Figure. 6.**
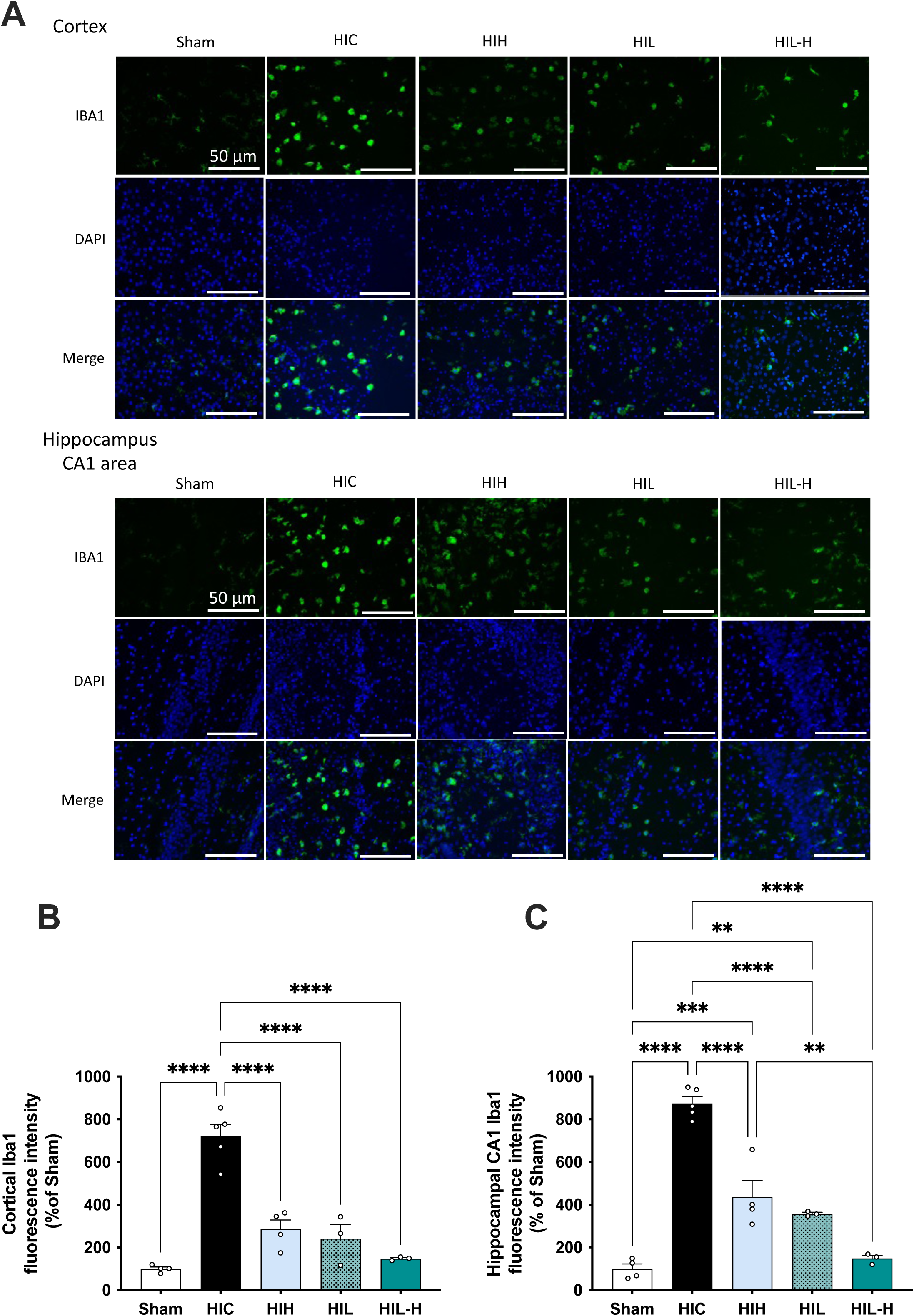
Early assessment of microglial activation. (A) Representative immunofluorescence images showing Iba1 immunoreactivity in the ipsilateral cortex and hippocampal CA1 region from Sham (n = 4), HIC (n = 5), HIH (n = 4), HIL (n = 3), and HIL-H (n = 3) groups 48h after NHI (P9). Images were acquired at 20× magnification. Scale bar, 50 μm. (B, C) Quantification of Iba1 fluorescence intensity in the cortex (B) and hippocampal CA1 region (C). For each animal, fluorescence intensity was averaged from two independent fields per brain region. Data are presented as mean ± SEM. Statistical analyses were performed using one-way ANOVA followed by Tukey’s multiple-comparisons test. Statistical comparisons were performed relative to the Sham group. Significance is indicated as **p < 0.01, ***p < 0.001, and **** < 0.0001.

### Western blot analyses of some key metabolic proteins following NHI

To investigate molecular mechanisms underlying lactate-mediated neuroprotection, cortical protein expression was analyzed by Western blot at P9 (**Fig.7**). In the ipsilateral hemisphere, expression of the neuronal lactate transporter MCT2 was reduced in HIC and HIH animals but remained comparable to Sham levels following lactate treatment, either alone or combined with TH (**Fig.7A**). Conversely, the expression of the astrocytic glutamate transporter EAAT1 was increased in HIC and HIH animals but was normalized in the HIL and HIL-H groups (**Fig.7B**). Finally, the expression of glucose-6-phosphate dehydrogenase (G6PDH), a key rate-limiting enzyme in the pentose phosphate pathway, was significantly higher in the HIL-H group than in the HIC and HIH groups, whereas HIL animals showed a similar trend that did not reach statistical significance (**Fig.7C**). No significant differences were detected in the contralateral cortex, except for G6PD, which was increased in the HIL group *vs.* Sham, HIC, and HIL-H groups (**Fig.S11A-C**).

**Figure. 7.**
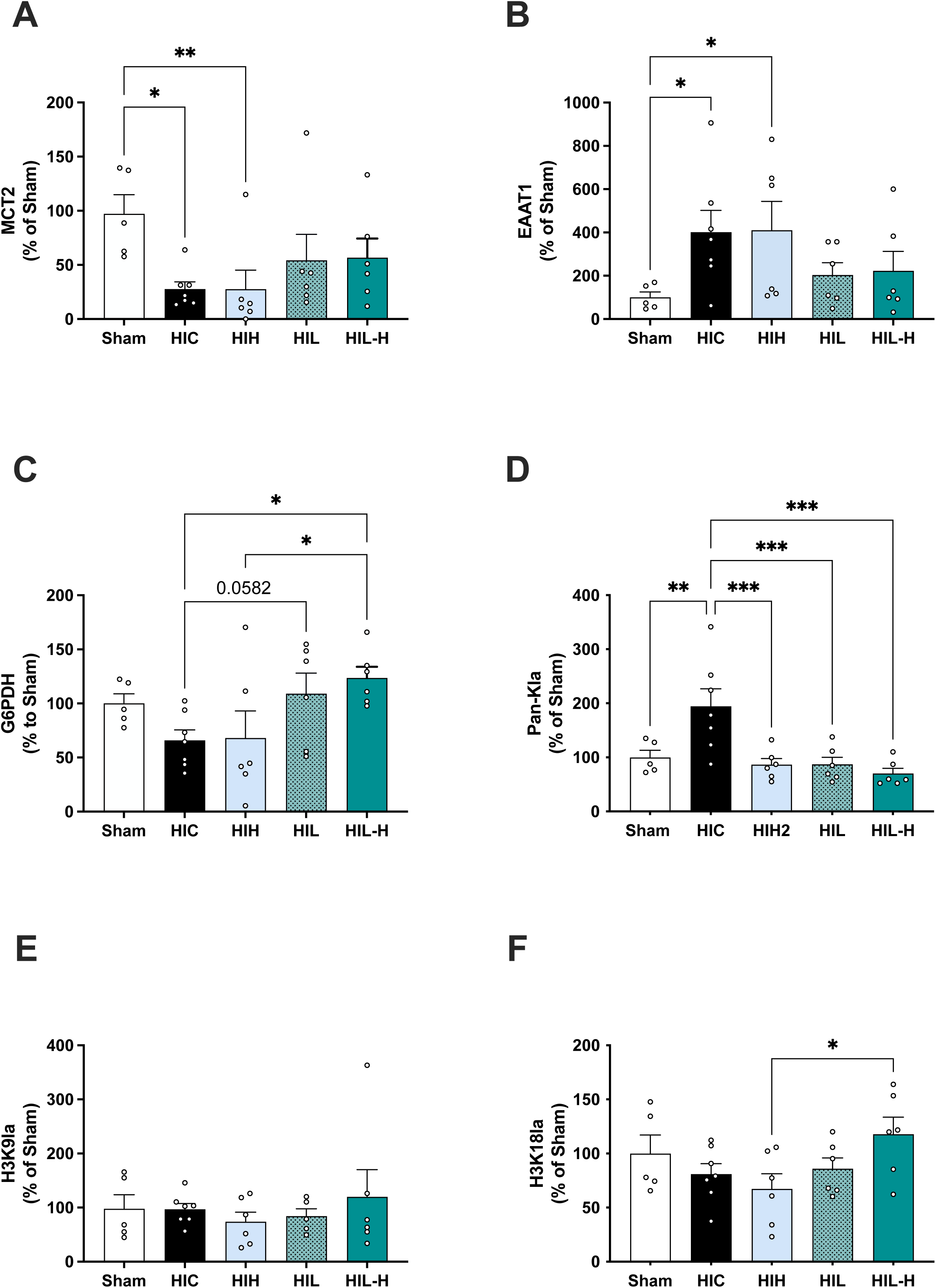
Modulation of metabolic protein expression and histone lactylation following NHI. (A–C) Western blot quantification of ipsilateral cortical expression of MCT2 (A), EAAT1 (B), and G6PDH (C) 48h after NHI (P9) in Sham (n = 5), HIC (n = 7), HIH (n = 6), HIL (n = 6), and HIL-H (n = 6) groups. (D–F) Quantification of cortical protein lactylation, including pan-protein lactylation (pan-Kla; D), H3K9 lactylation (H3K9la; E), and H3K18 lactylation (H3K18la; F), in the same experimental groups. Protein levels were normalized to total protein staining or β-actin, as appropriate, and expressed as a percentage of Sham values (100%). Data are presented as mean ± SEM. Statistical analyses were performed using ordinary one-way ANOVA followed by Fisher’s least significant difference (LSD) multiple-comparisons test for normally distributed data or Kruskal-Wallis tests followed by Dunn’s multiple-comparisons test for non-normally distributed data, as appropriate. Statistical significance is indicated as *p < 0.05, **p < 0.01, and ***p < 0.001.

### Lactylation following NHI

Pan-protein lysine lactylation (pan-Kla) was markedly increased in the ipsilateral cortex of HIC animals compared with all other groups (**Fig.7D**). Lactate, TH, and their combination prevented this increase, restoring pan-Kla levels to those of sham animals. No differences were detected in the contralateral hemisphere (**Fig.S11D**). Among site-specific histone lactylation marks, H3K9la remained unchanged across experimental groups (**Fig.7E**). In contrast, H3K18la tended to increase following lactate treatment and was significantly higher in the HIL-H group than in the HIH group in the ipsilateral hemisphere (**Fig.7F**). Similarly, no significant differences were detected in the contralateral cortex, except for H3K18la, which was increased in the HIL-H group relative to the HIC group (**Fig.S11D-F**).

### Time window for lactate administration

To examine the impact of lactate administration timing in combination with TH, animals receiving lactate before TH (HIL-H) were compared with those receiving the first lactate injection immediately after TH (HIH-L).

Longitudinal MRI demonstrated significantly smaller brain lesion volumes in the HIL-H group at both P9 (18 ± 2% vs. 30 ± 1%) and P30 (2 ± 2% vs. 21 ± 2%) (**Fig.8A and Fig.S12A**), indicating enhanced structural neuroprotection when lactate was administered before hypothermia. These MRI findings were accompanied by improved functional recovery. Compared with HIH-L animals, HIL-H animals exhibited better righting reflex performance at P12, lower mNSS scores at P24, and improved recognition memory at P45, with higher discrimination and recognition indices (**Fig.8B–D and Fig.S12B-C**). Early lactate administration also reduced latency to eat in the food restriction test, whereas bruxism did not differ significantly between groups (**Fig.8E–F**).

**Figure. 8.**
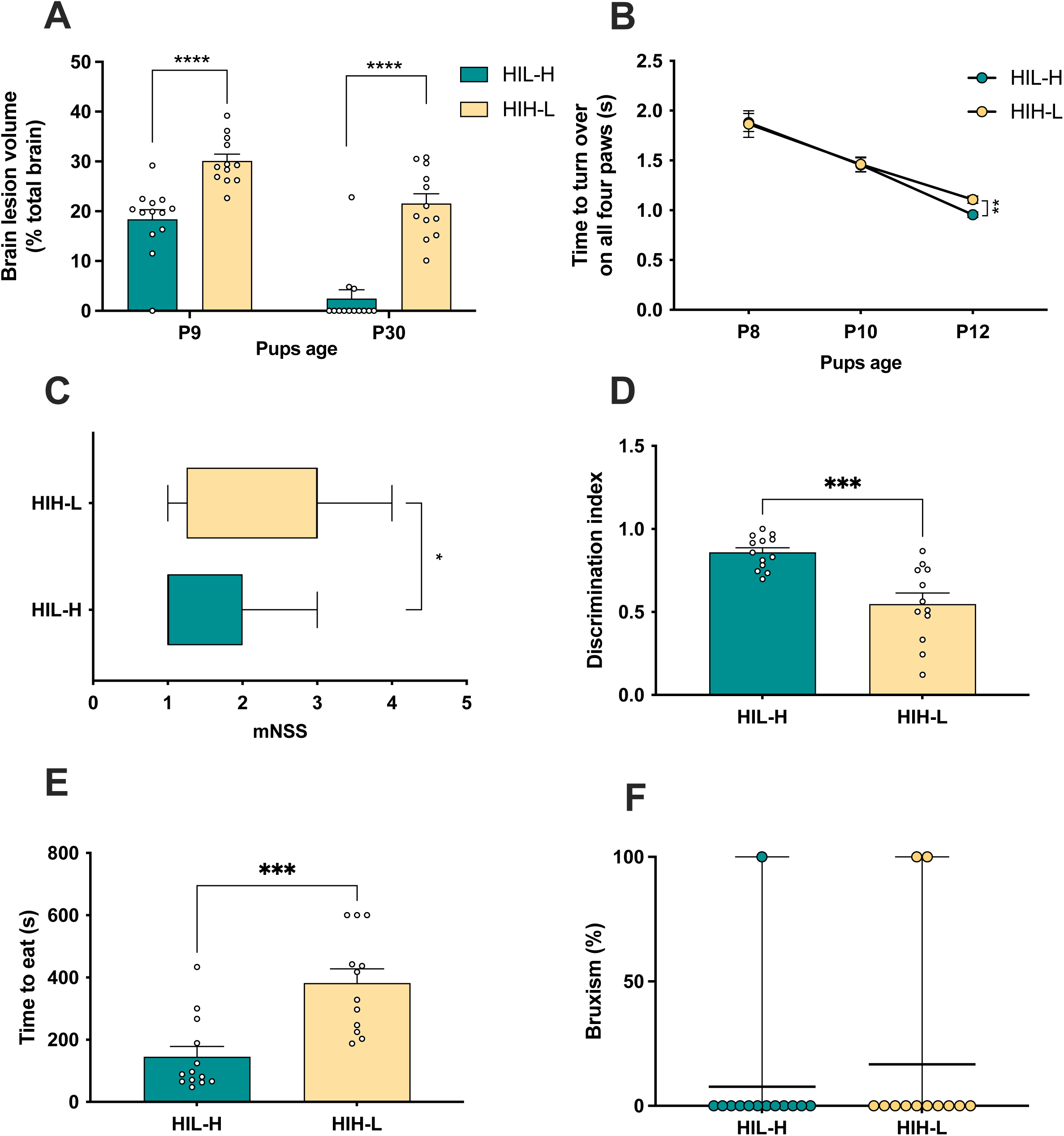
Impact of the time point of lactate administration. The first sodium L-lactate injection was administered either immediately after NHI and before the initiation of TH (HIL-H group) or immediately after completion of TH (HIH-L group). (A) Quantification of brain lesion volumes, expressed as a percentage of total brain volume. (B) Righting reflex assessed at P8, P10, and P12. (C) Modified Neurological Severity Score (mNSS) evaluated at P24. (D) Novel object recognition test performed at P45 showing the discrimination index. (E) Latency to consume the food pellet during the food restriction test performed at P51. (F) Proportion of animals exhibiting bruxism (%) during the food restriction test at P51. Analyses were performed in HIL-H (n = 13) and HIH-L (n = 12) groups. Data are presented as mean ± SEM, except for mNSS data, which are presented as median with interquartile range. Statistical analyses were performed using an unpaired Student’s t-test for normally distributed data or a Mann-Whitney test for non-normally distributed data, as appropriate. Statistical significance is indicated as *p < 0.05, **p < 0.01, ***p < 0.001, and ****p < 0.0001.

Together, these findings demonstrate that lactate administration before TH provides greater structural and functional neuroprotection than administration after TH, supporting lactate administration before the initiation of TH as the preferred therapeutic strategy for combined lactate-TH treatment in this experimental model.

## Discussion

This study demonstrates that lactate is neuroprotective following NHI and, importantly, that its neuroprotective effects are maintained when combined with TH, the current standard-of-care treatment for NHI. Using a multimodal longitudinal approach, we show that lactate administration, alone or in combination with hypothermia, reduced early lesion volume, attenuated apoptotic and inflammatory responses, preserved markers of metabolic adaptation, and improved long-term functional outcomes. Importantly, the combination of lactate and TH provided greater early structural and functional protection than TH alone, particularly when lactate was administered before the initiation of TH. These findings provide a strong rationale for considering lactate as an adjunctive therapy to TH in NHI.

### Lactate provides rapid neuroprotection following NHI

First, our findings confirm the rapid neuroprotective effect of exogenous lactate following NHI. A single administration of lactate immediately after NHI reduced MRI lesion volume by 26% within 30min. This finding is consistent with our previous study, in which lactate administration reduced lesion volume by 24% at the same early time point.^26^ Importantly, the present study used the commercially available sodium L-lactate formulation used in clinical practice. The dose administered was within the range used clinically with this formulation in very preterm infants for the correction of metabolic acidosis and has previously been shown to be well tolerated, without clinically relevant changes in blood lactate concentrations or acid-base status.^35^ Consistent with these clinical observations, lactate administration did not result in higher blood lactate concentrations in the present study compared with saline-treated animals (**Fig.S1**). During the kinetic analysis, blood lactate levels were even lower in lactate-treated animals at 180 and 300min after injection, suggesting rapid systemic clearance and/or metabolic utilization of the administered lactate. Thus, the present study uses a clinically available formulation and a dose range already established in the neonatal setting, further strengthening the translational relevance of our findings.^36^

The rapid metabolic utilization of exogenous lactate is further supported by the study of Roumes *et al.*, in which administration of ^13^C-labeled lactate resulted in the incorporation of ^13^C into brain amino acids as early as 30 min after administration.^26^ Moreover, it has been shown that co-administration of oxamate, an inhibitor of lactate dehydrogenase, abolished the neuroprotective effect of exogenous lactate and resulted in lesion volumes even greater than those observed in control animals.^26^ These findings suggest that endogenous lactate production during NHI may itself contribute to tissue recovery through its metabolic utilization, challenging the traditional view of lactate as merely a marker of metabolic distress or a metabolic waste product. This perspective calls for a re-evaluation of the role of lactate,^37^ in which lactate accumulation may reflect not only metabolic dysfunction but also an adaptive metabolic response aimed at sustaining energy production and promoting tissue recovery.

In addition to its direct use as an energetic substrate, lactate also modulate protein expression of key protein implicated in brain metabolism. Indeed, the neuronal lactate transporter MCT2 expression was reduced following NHI and remained decreased after TH alone, whereas lactate administration, either alone or combined with TH, preserved MCT2 expression and therefore its metabolic use. These results are consistent with previous studies showing that NHI and cerebral ischemia impair neuronal lactate transport capacity, while lactate supplementation helps preserve neuronal lactate transport under ischemic condition.^31,38^ Consistent with previous report, NHI also induced cortical EAAT1 (the astrocytic glutamate uptake transporter) upregulation, likely reflecting an adaptive response to glutamate accumulation and excitotoxic stress.^39^ Interestingly, this response was absent in lactate-treated animals, suggesting that lactate treatment may reduce the need for this compensatory mechanism response by limiting the glutamate excitotoxicity. Finally, we also observed an increase in G6PD expression in lactate-treated animals, potentially reflecting enhanced pentose phosphate pathway (PPP) activity and improved redox homeostasis, consistent with previous findings.^40^ The PPP is a major source of NADPH, which is required to maintain cellular antioxidant defenses and therefore represents an important metabolic pathway for limiting oxidative stress after HI. Because glucose is the primary substrate fueling the PPP, the increased reliance on lactate as an oxidative substrate may allow glucose to be preferentially directed toward the PPP rather than being used for energy production, thereby supporting redox homeostasis and contributing to neuroprotection. This result is further supported by data on apoptosis that was decreased in the lactate-treated groups (both caspase-3 labeling and Nissl staining), in line with previous reports.^26,41^

### Neuroprotective effects of lactate beyond its role as an energy substrate

Although these findings support a role for lactate as an energetic substrate, lactate may also exert neuroprotective effects through its function as a signaling molecule. Such signaling-dependent effects have been demonstrated in adult experimental stroke^22,42^ and have also been implicated in NHI,^33^ particularly for late recovery phenomenon. Thus, the neuroprotective effects of lactate are likely to reflect the combined contribution of its metabolic and signaling functions. In addition to its effect as a signal molecule through the receptor HCAR1, lactate may also have epigenetic function and modify gene expression, a phenomenon called lactylation. In this study, we observed different lactylation levels in the different groups. HI markedly increased pan-Kla levels in the ipsilateral cortex, whereas lactate, TH, and their combination restored pan-Kla to sham levels. Interestingly, H3K18la was selectively increased in lactate-treated groups, particularly following combined therapy, whereas H3K9la remained unchanged. These findings support the idea of lactylation-site-specific regulation. Selective H3K18la induction suggests a targeted epigenetic mechanism potentially involved in neuroprotection, as H3K18la has been suggested to limit neuroinflammation after ischemic stroke.^43,44^ Further studies are needed to determine whether H3K18 lactylation contributes to microglial functional remodeling and long-term neuroprotection following NHI and to identify downstream genes regulated by this modification.

Finally, we explored the impact of lactate on immunomodulation.^45^ Lactate alone and combined with TH reduced Iba1 immunoreactivity in the ipsilateral cortex and hippocampus 48h after NHI. lactate administration also reduced CD86, a marker associated with pro-inflammatory microglial activation, whereas combined treatment increased CD206, a marker associated with anti-inflammatory microglial activation,^46^ particularly in the hippocampus (**Fig.S10**). Together, Together, these findings suggest that lactate may limit the excessive pro-inflammatory response while promoting a more anti-inflammatory response. Further functional and transcriptomic studies are needed to determine whether this reflects sustained microglial reprogramming.

### Lactate neuroprotection is compatible with therapeutic hypothermia

The central finding of this study is that lactate neuroprotection is preserved when is preserved when combined with TH. This is particularly relevant clinically because TH is currently the only established neuroprotective treatment for NHI. Rather than interfering with the beneficial effects of TH, lactate administration provided additional neuroprotection, with the combined treatment producing the greatest early lesion recovery. Indeed, within the first 48h after NHI, lesion recovery reached approximately 51% in animals receiving combined lactate–TH treatment, compared with approximately 35% following lactate alone and 29% following TH alone. Although brain lesion volumes no longer differed significantly between treated groups at P30, the early structural advantage of combined treatment was accompanied by favorable functional outcomes. Importantly, MRI lesion volume does not necessarily fully reflect functional recovery, as similar degrees of structural injury may be associated with different behavioral outcomes.^47^ Longitudinal behavioral assessment is therefore essential to determine whether the structural protection observed by MRI translates into meaningful functional benefit. Concerning short-time behavior, animals in the HIL-H group exhibited complete righting reflex recovery by P12, lower mNSS scores at P24, and discrimination indices comparable to sham animals at P45, outperforming TH-treated animals across functional endpoints. Importantly, benefit of lactate administration before TH was superior to delayed administration, identifying an early therapeutic window consistent with observations for other adjunctive therapies, including melatonin^10,11^ and allopurinol.^48^

### Limitations of the study

This study has some limitations. Although no sex-related differences were detected across MRI and behavioral outcomes in the present study, consistent with some previous studies using the Rice-Vannucci model, findings across neonatal rodent studies remain inconsistent, with some reporting sex-dependent effects and others finding no significant differences.^34,49^ In contrast, sex-related differences have been more consistently reported in human neonatal HI.^50^ Dedicated studies in which sex is investigated as a primary variable will therefore be required to clarify whether and how sex influences the response to NHI and to neuroprotective interventions in neonatal rodents. In addition, *ex vivo* analyses were performed in a limited number of animals, as the study was primarily designed to enable longitudinal MRI and behavioral assessment while limiting animal use. Despite this limited sample size, significant differences between groups were detected across several histological, immunohistochemical, and protein expression analyses, indicating robust treatment-related effects. However, the restricted number of animals limited the extent of these analyses, with protein expression measurements mainly confined to the cortex at a single post-injury time point. Future studies should therefore investigate additional brain regions and multiple post-injury time points to better characterize the temporal and regional mechanisms underlying lactate neuroprotection combined to TH. Finally, validation of the combined lactate-TH strategy in clinically relevant large-animal models, such as neonatal piglets, would be an important additional step toward clinical translation.

## Conclusion

In summary, this study demonstrates that sodium L-lactate administration provides robust neuroprotection following NHI injury and, importantly, that its effects are preserved and even enhanced when combined with therapeutic hypothermia. Our recent clinical study further showed that sodium L-lactate infusion was well tolerated in preterm infants, supporting its potential for clinical translation.^35^ Together, these findings provide a rationale for further evaluating sodium L-lactate as an adjunct to hypothermia in newborns with hypoxic-ischemic encephalopathy.

## Acknowledgements

This study was conducted within the framework of the University of Bordeaux IdEx Investments for the Future program (RRI IMPACT).

## Funding

This work was supported by the French National Research Agency (ANR) through the BrainFuel project (grant ANR-21-CE44-0023 to A.K.B.S. and L.P.), the SPARK program, the Fédération pour la Recherche sur le Cerveau (FRC), and the Région Nouvelle-Aquitaine.

## Disclosures

The authors declare no competing interests.

## Notes

### Competing Interest Statement

The authors have declared no competing interest.

